# Biofilm thickness matters: Deterministic assembly of different functions and communities in nitrifying biofilms

**DOI:** 10.1101/416701

**Authors:** Carolina Suarez, Maria Piculell, Oskar Modin, Silke Langenheder, Frank Persson, Malte Hermansson

## Abstract

Microbial biofilms are important in natural ecosystems and in biotechnological applications. Biofilm architecture influences organisms’ spatial positions, who their neighbors are, and redox gradients, which in turn determine functions. We ask if and how biofilm thickness influences community composition, architecture and functions. But biofilm thickness cannot easily be isolated from external environmental factors. We designed a metacommunity system in a wastewater treatment plant, where either 50 or 400 µm thick nitrifying biofilms were grown simultaneously on biofilm carriers in the same reactor. Model simulations showed that the 50 µm biofilms could be fully oxygenated whereas the 400 µm biofilms contained anaerobic zones. The 50 and 400 µm biofilms developed significantly different communities. due to deterministic factors were stronger than homogenizing dispersal forces in the reactor, despite the fact that biofilms experienced the same history and external conditions. Relative abundance of aerobic nitrifiers was higher in the 50 µm biofilms, while anaerobic ammonium oxidizers were more abundant in the 400 µm biofilms. However, turnover was larger than the nestedness component of between-group beta-diversity, i.e. the 50 µm biofilm was not just a subset of the thicker 400 µm biofilm with reduced taxa richness. Furthermore, the communities had different nitrogen transformation rates. The study shows that biofilm thickness has a strong impact on community composition and ecosystem function, which has implications for biotechnological applications, and for our general understanding of biofilms.

**IMPORTANCE:** Microorganisms colonize all surfaces in water and form biofilms. Diffusion limitations form steep gradients of energy and nutrient sources from the water phase into the deeper biofilm parts, influencing community composition through the biofilm. Thickness of the biofilm will affect diffusion gradients, and is therefore presumably important for biofilm composition. Since environmental factors determine thickness, studies of how thickness influences biofilm functions and community assembly, have been difficult to perform. We studied biofilms for wastewater treatment with fixed thicknesses of 50 and 400 µm during otherwise similar conditions and history. Despite growing in the same wastewater reactor, 16S rRNA gene sequencing and confocal microscopy showed the formation of two different communities, performing different ecosystem functions. Using statistical methods, we show for the first time, how biofilm thickness influences community assembly. The results help our understanding of the ecology of microbial biofilms, and in designing engineered systems based on ecological principles.

## INTRODUCTION

Biofilms are dense communities, encased in a polymer matrix, attached to a surface and/or each other (1) with a high microbial diversity compared to the bulk water system (1-3). Microbial biofilms are important in aquatic ecosystems and are useful in many biotechnological applications, such as wastewater- or drinking water treatment. In nitrogen removal from wastewater, moving bed biofilm reactors (MBBRs) are used at many wastewater treatment plants (WWTPs). Here biofilms grow on so-called carriers, which move freely in the bioreactor (Fig. 1C). An MBBR can be seen as a metacommunity (4), just as other WWTP systems such as activated sludge flocs and granules, where each free-floating biofilm carrier represents a local community. The local communities have defined boundaries and are separate, but linked by dispersal with all other biofilm carriers in the reactor, in this case fed with wastewater from a full-scale WWTP to form nitrifying biofilms. The reactor represents the regional level and is assumed to be a spatially implicit system, i.e. dispersal is likely to be equal between all carriers (4).

**Fig. 1.**
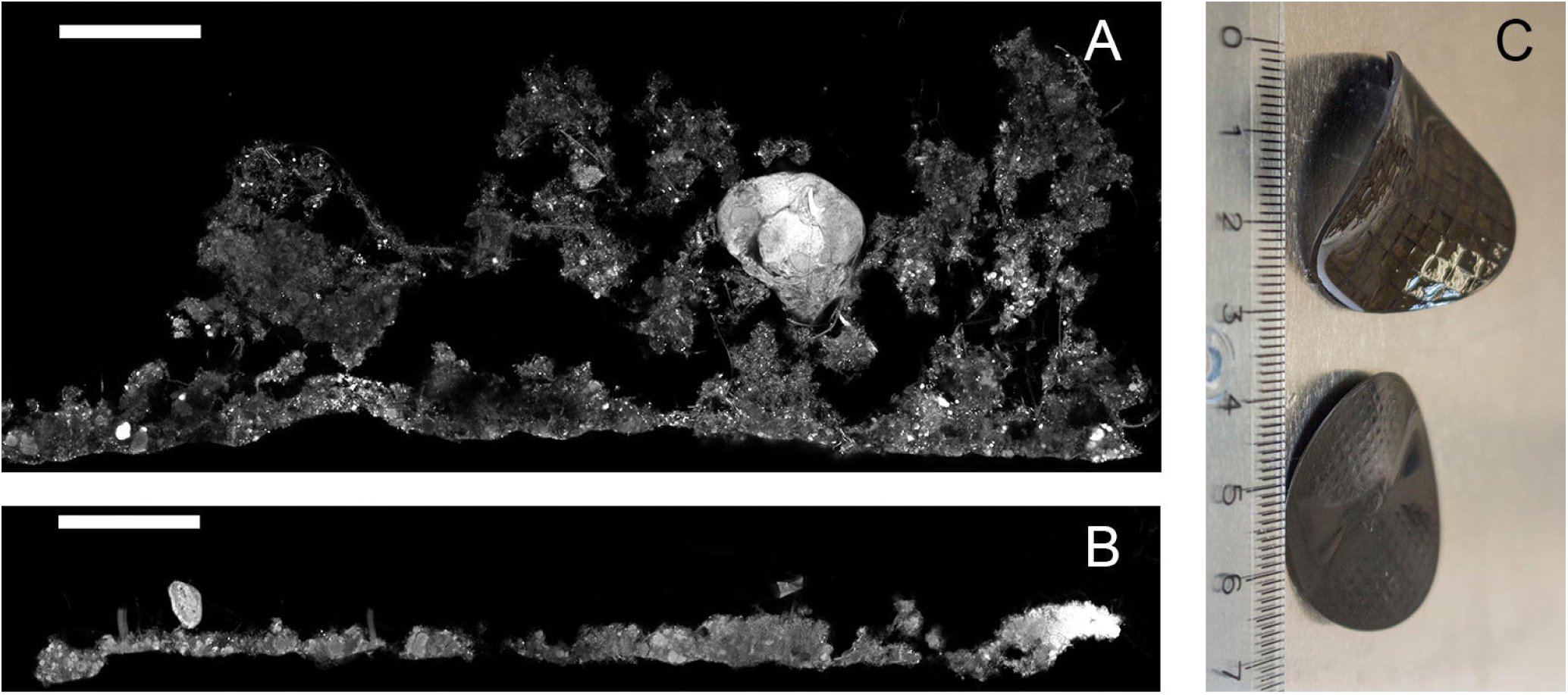
Biofilm structure shown by EPS staining of cryosections. The biofilm-water interface is the upper side. A. Z400 biofilm. B: Z50 biofilm. Scale bar: 100 µm. (C): Z400 (up) and Z50 (down) biofilm carriers; a ruler in cm is shown for size comparison.

The mechanisms of community assembly are central in microbial ecology and in our understanding of formation of biodiversity in all ecosystems (5, 6), including microbial communities in biofilms (7). Communities are formed by deterministic and stochastic factors that include four major ecological processes: selection, drift, dispersal and speciation (8). Selection, i.e. the sorting of species by prevailing local abiotic and biotic conditions, is deterministic, while drift results from stochastic birth and death events (6, 8). If local communities are further exposed to stochastic dispersal from the regional species pool (as we can assume to be the case of bioreactors in WWTPs), the expected result is that the abundance of a taxon in a local community can be predicted based on its respective abundance in the regional species pool and thereby follows neutral distribution patterns (9, 10). However, dispersal can also be deterministic if microorganisms differ in their ability to disperse within the complex spatial biofilm matrix or if their propagation is affected by interactions with species already present in the biofilm.

Selection has been suggested as the major mechanism for community assembly in stream biofilms (11-13), and other biofilms (14), while for biofilms within lakes linked by dispersal, both stochastic and deterministic factors were shown to be important (15). The importance of both stochastic and deterministic factors was shown in an elegant study using parallel microbial electrolysis cells incubated with wastewater (16). Other studies in wastewater activated sludge systems have shown the importance of deterministic and stochastic factors (17-20).

Deterministic assembly in biofilms could be due to specific mechanisms: Firstly, diffusion limitations form steep gradients of electron donors and acceptors in biofilms, which result in structured micro-environments. Examples are found in biofilms in WWTPs used for nitrification (i.e. oxidation of ammonium to nitrate) during the nitrogen removal process (21, 22). Here population stratification typically occurs; ammonia oxidizing bacteria (AOB) are found closest to the oxygenated water and nitrite oxidizing bacteria (NOB) below the AOB (23-26). If oxygen is consumed, anaerobic ammonium oxidizing (anammox) bacteria can establish in the deeper parts of the biofilm (23, 27, 28). Similarly, in other multispecies biofilms anaerobic sulfate reducing bacteria are found in the biofilm interior (29). However, functions in microbial communities are not always sorted according to such a thermodynamic “redox tower” of electron acceptor (30). This makes detailed *a priori* predictions of community structure in biofilms difficult. Secondly, in addition to gradients, it was realized early on that microbial biofilms are in fact complex structures and not homogenous layers of randomly distributed organisms (3) and, ever since, architecture has been viewed as an important biofilm property. The intricate biofilm architecture consists of towers, mushroom-like structures and water filled channels (1, 2, 7, 31). If biofilms differ in their architecture, dispersal effects could influence community assembly by changing the colonization surface available. Furthermore, microorganism with deterministic dispersal might show preference towards different types of biofilms.

Thickness will influence biofilm architecture and redox gradients and thus generally the local biofilm environment. However, the experimental evidence for effects of thickness on architecture, community structure and function has been difficult to obtain because biofilm thickness is the result of environmental conditions such as flow (1, 7, 32-34), nutrient conditions (24), development age of the biofilm (34), carbon to nitrogen (C/N) ratios (25) and temperature (35). In most experimental systems, thickness cannot easily be isolated from these environmental factors that themselves can influence the community structure and functions.

Recently, a biofilm carrier with a defined grid wall height that defines the biofilm thickness was designed (Z-carriers, Veolia Water Technologies AB – AnoxKaldnes, Lund, Sweden) (36-38). These carriers allow stringent experiments with different biofilm thicknesses, which have shown that thickness can affect some biofilm functions (38, 39), evenness (39), biofilm architecture (37), abundance of key organisms (37, 39) and functional stability after a disturbance (37). Beside the opportunities to gain basic ecological knowledge by designing new experiments, the ability to control biofilm thickness opens for new process configurations in WWTPs. In this study, a pilot nitrifying bioreactor was filled with a mixture of Z-carriers with biofilm thicknesses of 50 and 400 µm. Thus, environmental conditions and history of the biofilms were the same. We ask if thickness, in itself or via diffusion effects or other mechanisms, is important for bacterial community structure, and if so, what the possible mechanisms of community assembly would be. The thicknesses we investigated are within the range commonly found in natural- as well as in man-made biofilms (25, 26, 32, 40, 41).

Differences in communities, i.e. beta-diversity between thin and thick biofilms, could arise due to (a) turnover (species replacement) or (b) nestedness resulting from differences in alpha-diversity (42), which can both result from deterministic (e.g. environmental gradients) and stochastic factors (e.g. dispersal and drift) (42, 43). For example, as thicker biofilms have larger volume and surface-area, they are by chance expected to be colonized by more species, leading to higher richness. A possible deterministic mechanism for higher richness in 400 compared to 50 µm biofilms would be if redox profiles differ between them. Generally, the nitrifying bioreactor is an oxic environment and the presence of aerobic taxa is expected. However, if anoxic regions exist in thick biofilms, this would allow the establishment of anaerobic taxa, increasing species richness. In this case, we hypothesize that the 50 µm biofilm community would be a subset of the richer 400 µm biofilm community, due to anaerobic taxa being restricted to the thicker 400 µm biofilms, while the same aerobic taxa occur in both the 50 and 400 µm biofilms. Alternatively, turnover, i.e. differences in species identity, could arise if biofilms of different thickness have different environmental conditions apart from redox profiles. Finally, we also measured rates of nitrogen transformations in the two biofilms and investigated whether possible differences in ammonia- and nitrite removal rates between biofilms of distinct thicknesses are primarily linked to differences in taxa richness (i.e. nestedness) or identity (i.e. turnover) and discuss the implications of the results for wastewater treatment.

## RESULTS

### Two different biofilms

We grew biofilm communities with a maximum thickness of 50 μm and 400 μm together in the same bioreactor; these communities are referred to as Z50 and Z400. CLSM images of EPS stained biofilm cryosections confirmed that carrier design limited biofilm thickness (Fig. 1A and B).

### Alpha and beta diversity

The alpha diversity parameters richness (^0^D), first-order diversity (^1^D) and evenness (^1^D/^0^D) (Fig. 2A), were all significantly higher for the thick Z400 biofilms than for the thinner Z50 biofilms (Welch t-test, p<0.05). We also estimated beta diversity using the presence-absence based Sørensen index (β_sor_), which showed that Z50 and Z400 communities were different (PERMANOVA β_sor_, p=0.002, r^2^=0.50) (Fig. 2B).

**Fig. 2.**
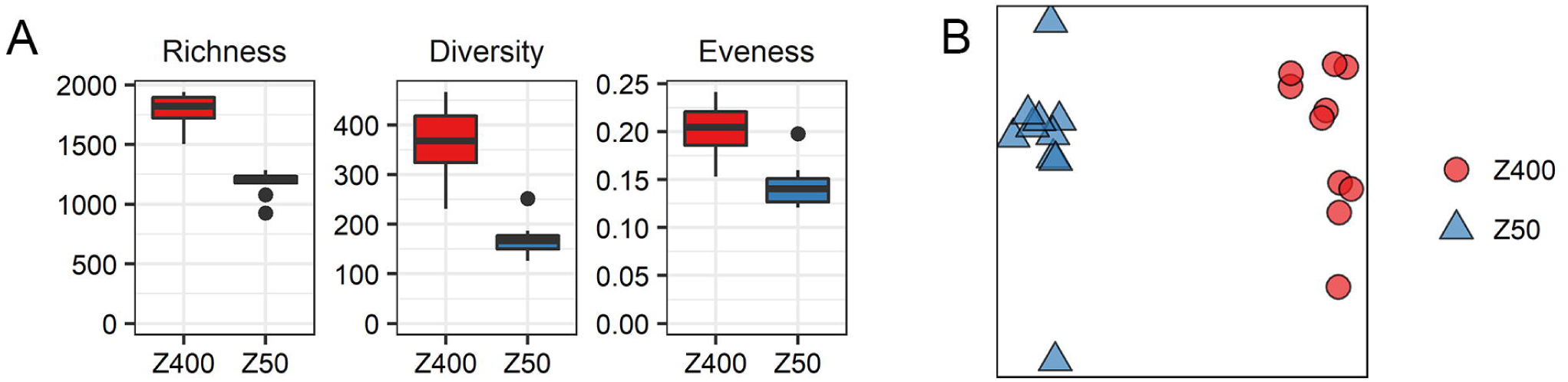
A: Richness (^0^D), diversity (^1^D) and evenness (^1^D/ ^0^D) for the Z50 and Z400 biofilms. B: PCoA based on the Sørensen index (β_sor_).

We used null modelling to estimate the standardized effect size (SES) for β_sor_. We observed that β_sor_ values for between-group comparisons, i.e. between Z50 and Z400, were higher than expected by chance β_sor_ values for between-group comparisons (SES_βsor_ > +2) (Fig. 3A) indicating that between-group differences were likely deterministic. On the contrary, observed β_sor_ values for within-group comparisons, i.e. between the carriers of the same type, were not more different than expected by chance (**|**SES_βsor_**|** < 2) (Fig. 3A). In addition, estimation of the quantitative RCBRAY metric, also indicated that Z50 and Z400 communities were in average more dissimilar than the null expectation (between-group RCBRAY > +0.95)

**Fig. 3.**
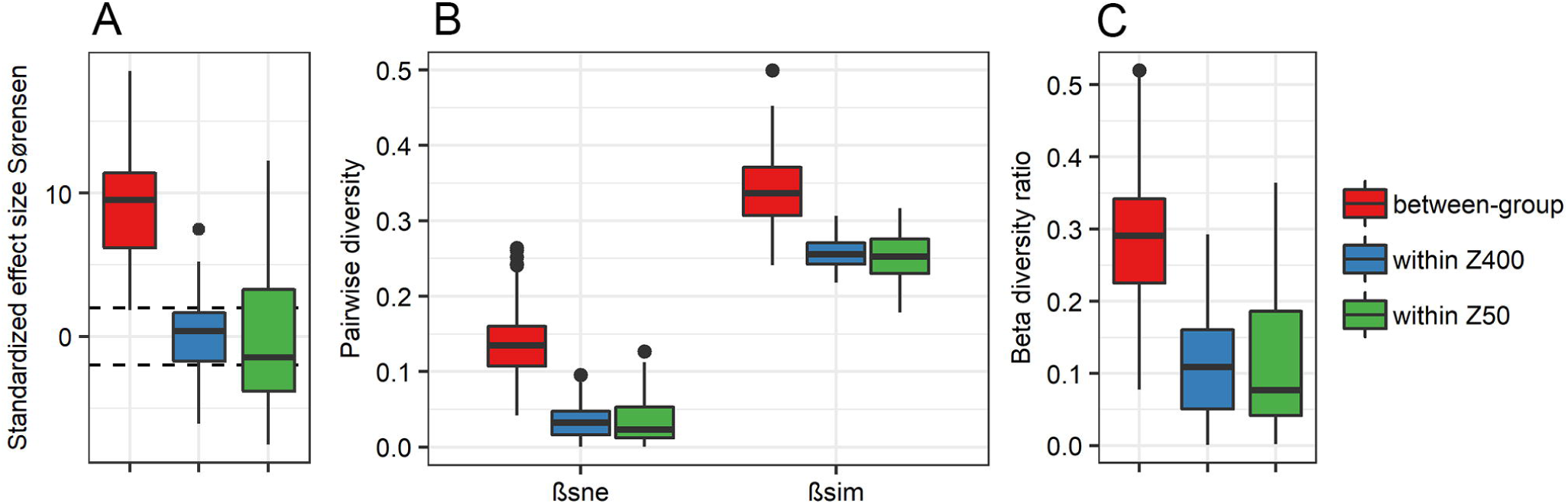
(A) Standardized effect size for the Sørensen index (β_sor_); dashed lines indicate SES values of +2 and −2. B: β_sne_ (dissimilarity due to nestedness) and β_sim_ (turnover) values; the sum of β_sim_ and β_sne_ is β_sor_. (C) Beta diversity ratio. Values were estimated for pairwise comparisons among Z400 replicates (n=10), Z50 replicates (n=10) and between the two groups.

To determine whether the between-group beta diversity was due to nestedness or turnover, we estimated the components of β_sor_; turnover (β_sim_) and dissimilarity due to nestedness (β_sne_) using the Baselga framework (42) (Fig. 3B), to estimate the β_ratio_ (44). When β_ratio_ is smaller than 0.5, beta diversity is dominated by turnover rather than β_sne_ (44). We observed a between-group β_ratio_ lower than 0.5 (Fig. 3C). The differences between the Z50 and Z400 communities due to turnover were significant (PERMANOVA β_sim_, p=0.001, r^2^=0.34). Thus, the beta diversity between the Z50 and Z400 communities was caused by both nestedness and turnover, with turnover being more important.

Differences in relative abundance of taxa between Z50 and Z400 were estimated using DESeq2. We found differential abundance (p_(adj)_<0.01, DESeq2) for 45% of the sequence variants (SVs) analyzed with DESeq, while for the top 40 most abundant SVs, 32 had different abundance between Z50 and Z400 (Fig. S1). Among the fraction with differential abundance, 26% of SVs were more abundant in Z50, and 74% were more abundant in Z400. The effect of thickness on relative abundance, if any, differed among taxa (for example see Fig. S1).

### Between-group sorting of nitrifiers and anammox bacteria

The relative read abundance of the nitrifiers, *Nitrosomonas, Nitrospira* and *Nitrotoga*, was lower in the Z400 biofilms with *Nitrotoga* being almost restricted to Z50 (Fig. 4A). The same trends were noticed using quantitative fluorescence in situ hybridization (qFISH; Fig. 4C; Welch’s t-test, p<0.05). It was not possible to detect by qPCR if comammox were present due to non-specific amplification using *Nitrospira amoA* primers (45).

**Fig. 4.**
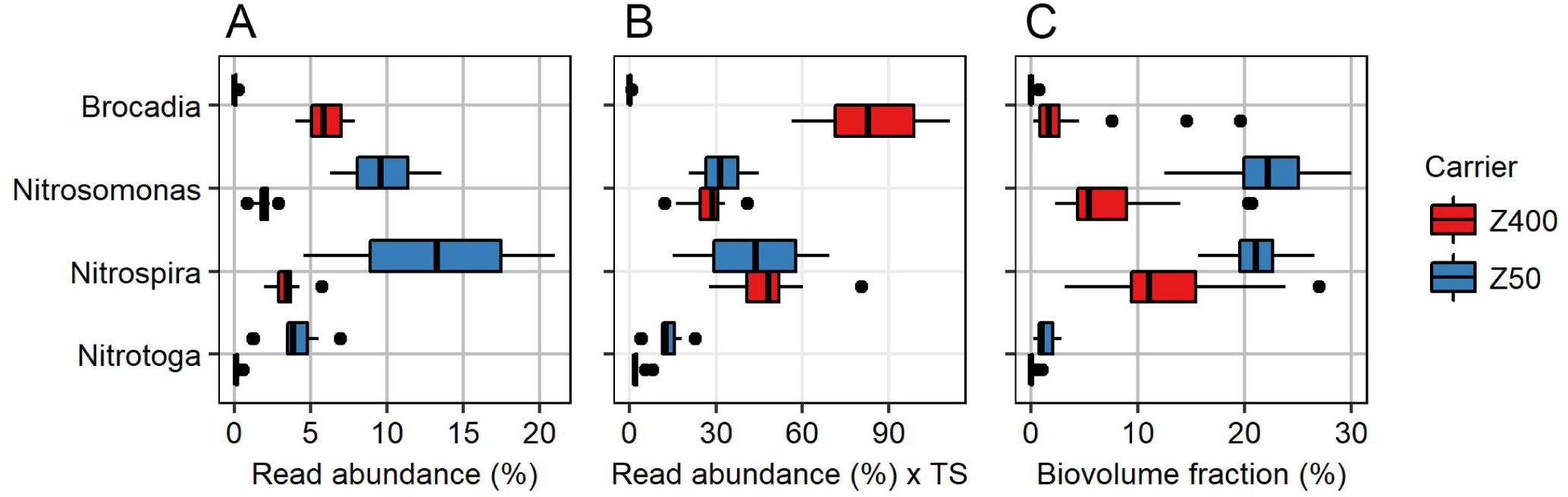
(A) Relative read abundance of nitrifiers and anammox bacteria in Z50 and Z400. (B) Relative read abundance multiplied by total solids (TS) measurements for each carrier type. (C) Biovolume fractions of nitrifiers and anammox bacteria, as measured by qFISH.

Interestingly, we observed the anammox bacterium *Brocadia* in the Z400 biofilms, but it was almost absent in the Z50 biofilms (Fig. 4A). This was supported by qFISH (Welch t-test p<0.001) (Fig. 4C). Sorting of bacteria between thick and thin biofilms was not only limited to primary producers (i.e. autotrophic nitrogen converters) but also seen among the predatory *Bdellovibrionales*. *Bacteriovorax* had a higher abundance in the Z50 communities, while some SVs classified as *Bdellovibrio* were more abundant in either Z400 or Z50 (Fig. S2).

FISH analyses of biofilm cryosections showed that the Z400 biofilm was stratified, e.g. with *Nitrospira* being more abundant in the middle of the biofilm and the anaerobic anammox bacteria being present in the deeper layers; a lack of stratification was observed *Nitrosomonas* (Fig. 5A and 5B). In the thin Z50 biofilms, no stratification was observed as the AOB and NOB populations were located side by side (Fig. 5C). The calculated dissolved oxygen (DO) concentration profiles in the biofilms are shown in Fig. 6; the results give a range of possible DO concentration profiles, which are shown as shaded regions. The model predicts that Z50 biofilms can be fully oxygenated but may also have anoxic regions, whereas the Z400 biofilms contain a completely anoxic region in its deeper parts in all tested scenarios.

**Fig. 5.**
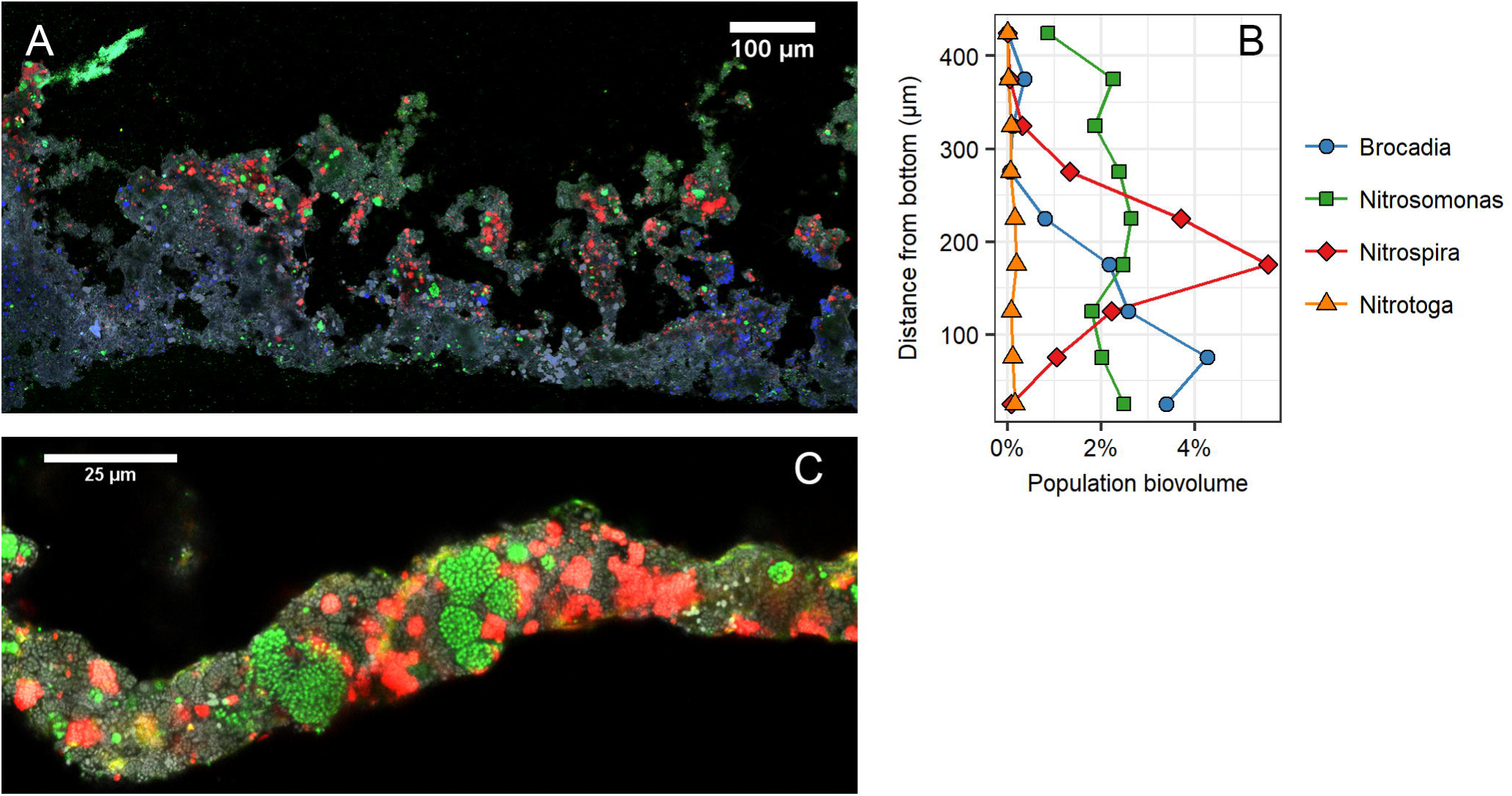
(A) FISH image of a Z400 biofilm cryosection; the water-biofilm interface is on the top. Green: *Nitrosomonas*. Red: *Nitrospira*. Yellow: *Nitrotoga*. Blue: *Brocadia*. Grey: SYTO. (B) FISH-based population distribution at different biofilm depths in Z400. (C) FISH image of a Z50 biofilm cryosection; the water-biofilm interface is on the top. Green: *Nitrosomonas*. Red: *Nitrospira*. Yellow: *Nitrotoga*. Grey: SYTO.

**Fig. 6.**
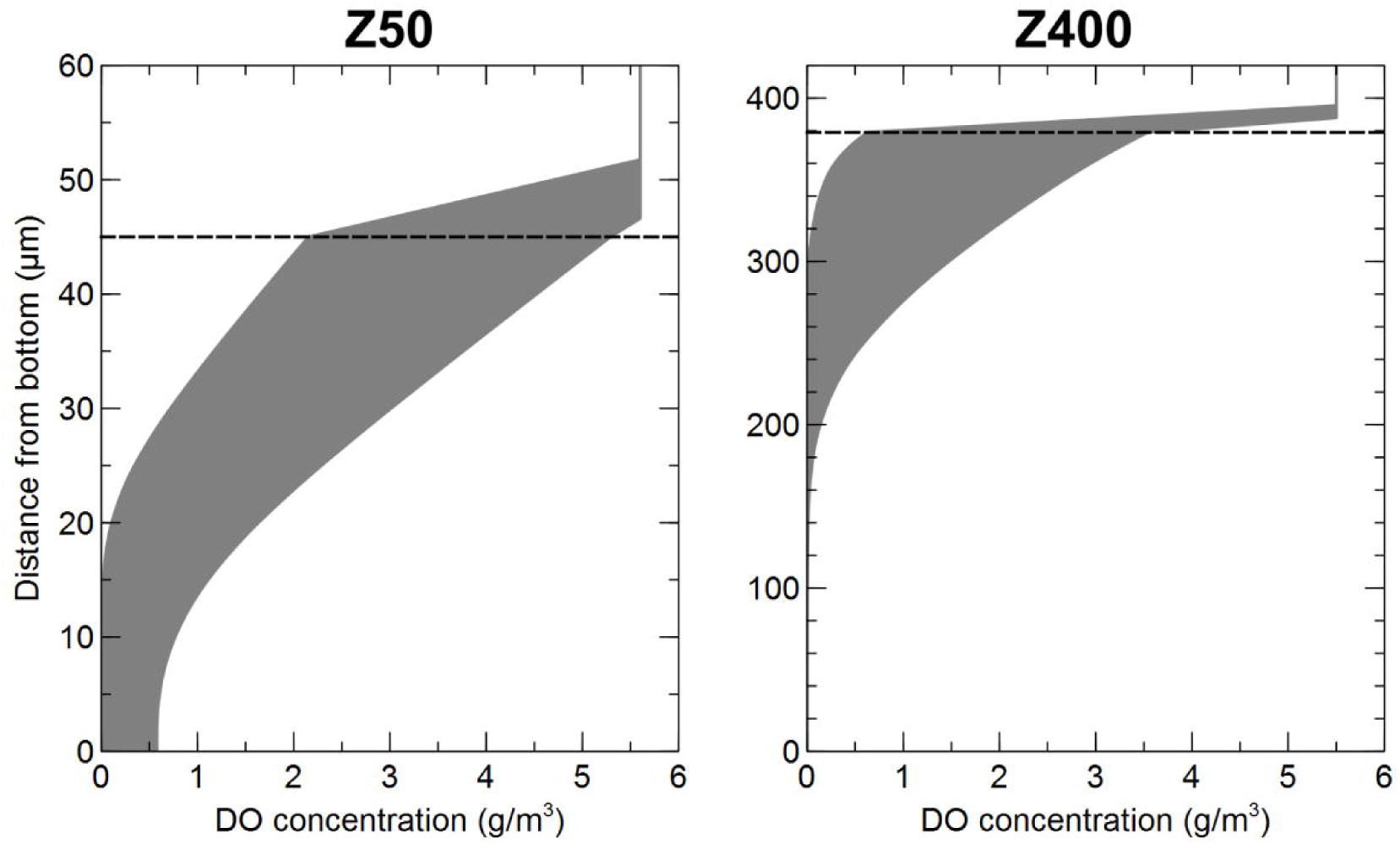
DO concentrations profiles in the Z50 and Z400 biofilms. The shaded regions show ranges of DO concentration profiles resulting from different assumption about the fraction of the total dry solids on the carriers that is active bacteria. The dashed horizontal lines show the biofilm-liquid interface.

### Nitrogen transformation rates

Two types of tests were performed separately on the Z50 and Z400 carriers; (i) actual activities tested in a continuous laboratory trial, with the same incoming water as in the 0.5 m^3^ reactor and (ii) potential activities tested in batch trials where excess nitrogen was added. For all trials, removal rates are reported per surface area and day. Actual rates of net NO_3_^-^ production were 1.4-1.5 gNO_3_^-^-N/m^2^, d for Z50 and 0.68-0.72 gNO_3_^-^-/m^2^, d for Z400. To estimate NO_3_^-^ production per nitrifier abundance, it is necessary to consider differences in biomass between carriers. We estimate that the total nitrifier biomass per carrier surface was about the same in Z50 and Z400 (Fig. 4B). Therefore, per nitrifier biomass, net NO_3_^-^ production was higher in Z50 than in Z400.

In the aerobic potential tests for net NH4+ removal (Fig. 7A), net NO_3_^-^ and net NO_2_^-^production (per carrier area) was higher for Z50 than Z400 biofilms (ANCOVA, p<0.05), while the rate of net NH4+ removal was not significantly different between Z50 and Z400 (ANCOVA, p>0.05). The aerobic potential removal of NO_2_^-^ (Fig. 7B) was significantly higher for Z400 than for Z50 (ANCOVA, p<0.05). Finally, in the anoxic potential trials, in which NH4+ and NO_2_^-^ were added simultaneously (Fig. 7C), removal of NO_2_^-^ was significantly higher for Z400 than for Z50 (ANCOVA, p<0.05), while no significant removal or production of NH4+ was seen for either Z50 or Z400.

**Fig. 7.**
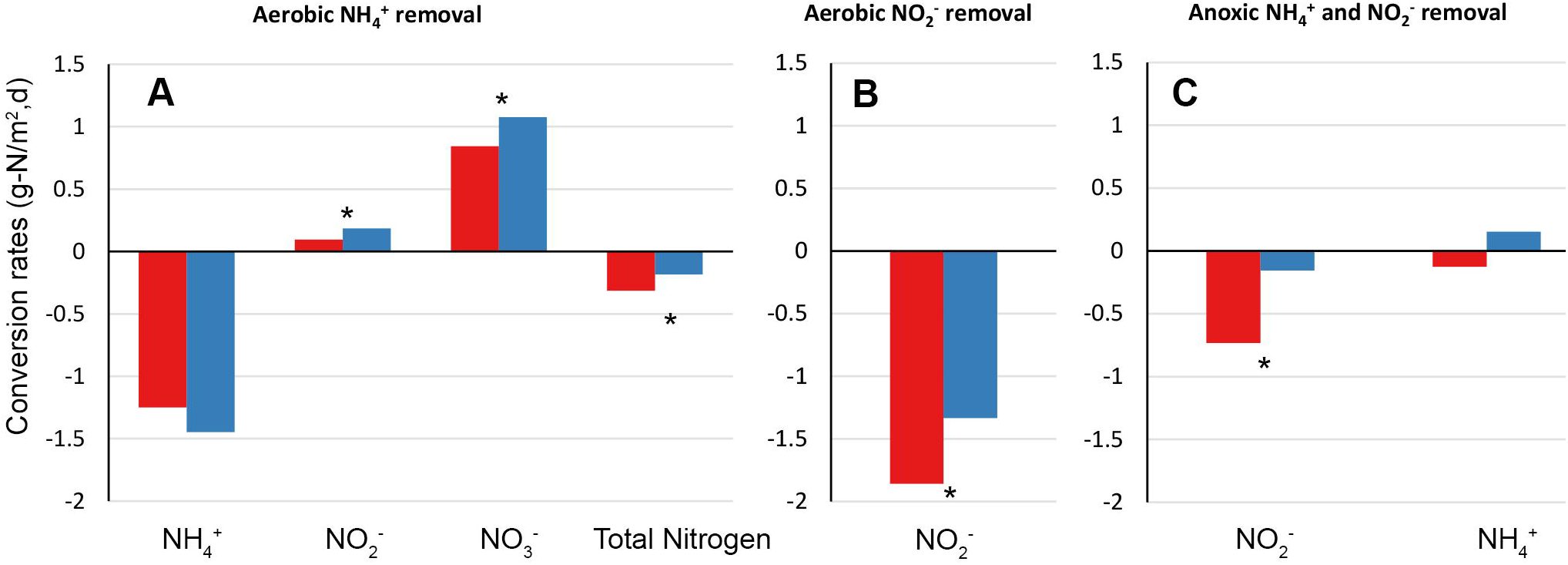
Potential conversion rates by carrier type during aerobic oxidation of NH4+ (A), aerobic oxidation of NO_2_- (B) and anoxic oxidation of NH4 (C) during batch tests. Significant differences between Z50 and Z400 (ANCOVA, p<0.05) are shown with (*). Red: Z400, Blue: Z50.

## DISCUSSION

Although incubated in the same bioreactor and experiencing the same conditions and the same history, different microbial communities developed on carriers with thin and thick biofilms (Fig. 2B). The thicker Z400 biofilm had a higher richness and evenness than the thinner Z50 biofilm (Fig. 2A) and our results are therefore in agreement with known positive species-area relationships for microbial communities (46). Moreover, similar to our results, Torresi et al. (39), focusing on micro-pollutant degradation, also found a significant higher evenness in thicker biofilms.

A null model approach was used to investigate if the differences in beta-diversity between Z50 and Z400 were due to deterministic or stochastic factors while accounting for the large differences in richness between Z50 and Z400 (47). The results showed that the between-group beta-diversity was higher than expected by chance (Fig. 3A), suggesting deterministic assembly due to differences in biofilm thickness. This result was also confirmed by the fact that biofilm thickness significantly affected the relative abundance of the majority of the most abundant individual taxa, meaning that they showed clear preference for either thin or thick biofilms (Fig. S1). Some turnover among the Z50 and Z400 replicates was observed, and was also expected due to ecological drift. Low SES values (Fig. 3A) suggest stochastic assembly among replicates, however, the relative importance of drift and dispersal cannot be disentangled with the experimental setup used here. In addition, due to the limited number of within-group replicates, these results should be interpreted with caution. Because MBBRs allow a high level of replication in communities linked by dispersal, a similar setup to the one use here with higher replication could be used to study stochastic assembly and to confirm the possible existence of alternate states (16, 48). Overall, other studies have shown that stochastic and deterministic processes can co-occur in biofilms (15, 17, 18). Our results suggest that the importance of deterministic vs. stochastic assembly depends on biofilm thickness: assembly would be deterministic between biofilms of different thickness, while assembly would likely be stochastic among biofilms with the same thickness.

Our hypothesis was that the communities in Z50 would be an aerobic subset of the ones in Z400. Thus, beta-diversity between Z50 and Z400 would largely be due to nestedness, whereas turnover would have a small contribution. This was expected due to different redox profiles between Z50 and Z400 biofilms (Fig. 6) which could create nestedness; oxygen in the thin Z50 biofilm inhibit the growth of obligate anaerobes like anammox bacteria (49). Thus, richness in Z400 would be higher, because the community is a mixture of aerobic and anaerobic taxa. Surprisingly, although between-group β_sne_ was observed, the β_ratio_ was below 0.5 (Fig. 3C), indicating that beta-diversity was dominated by turnover. Thus, the Z50 biofilm was not just a subset of the oxic upper layers of the Z400 biofilm, but differences were primarily due to turnover of taxa. For example, *Nitrotoga* was observed in Z50, but was nearly absent in Z400 (Fig, 4, S1), which cannot be easily explained by redox profiles. Independently of the mechanism, it appears that thin biofilms favor the NOB *Nitrotoga*, which could have consequences for operational strategies in wastewater treatment.

Redox profiles (Fig. 6) explain the stratification of some taxa like anammox bacteria and *Nitrospira* in the Z400 biofilm (Fig. 5B). *Nitrosomonas* was the dominant population at the top of the Z400 biofilm (Fig. 5A, S3C) and was also abundant in Z50. However, *Nitrosomonas* aggregates were present throughout the Z400 biofilm, even in regions predicted to be anoxic (Fig. 5A, 5B). Furthermore, in the thin Z50 biofilm, *Nitrospira* was seen alongside *Nitrosomonas* (Fig. 5C), and here its relative abundance was actually higher than in Z400. Hence redox profiles alone cannot explain the distribution of taxa in the reactor. The fact that redox is not the only determinant of the distribution of microorganisms, even in strongly structured environments like sediments, has been noted (30). The Z50 and Z400 biofilms also differed in their spatial structure, with Z50 being more dense and having a smoother architecture, compared with the Z400 (Fig. 1) (37); furthermore extracellular nucleic acids were observed in Z400 but not in Z50 (data not shown). Thus, these differences could contribute to the deterministic turnover observed in this study, by either selection or deterministic dispersal. Another possible mechanism for between-group species turnover are biotic interactions. For instance, some SVs within the predatory *Bdellovibrionales* were differently distributed between the biofilms (Fig. S2). It is plausible that the two biofilms represented different prey communities that in turn shaped the predatory *Bdellovibrionales* communities. Such influence on the predatory *Bacteriovorax* has been shown, even for closely related preys (50, 51). Furthermore, Torsvik et al. suggested that predation can act as a major factor driving prokaryotic diversity (52). Hence, biological interactions, such as predation, could have had a large effect on these biofilm communities, as shown for other wastewater biofilms (53).

Differences in community composition between Z50 and Z400 were to a larger extent determined by turnover than nestedness. Therefore, we predict that differences in nitrogen transformation rates among them might not necessarily be linked to the differences in richness and evenness between Z50 and Z400. This is despite previous examples that have shown that species richness (46) and evenness (54) may by themselves lead to higher productivity. Similar to earlier studies (39), we found that the thinner biofilm had higher net NO_3_^-^ -production rates, despite having lower richness. This supports that species composition might be more important than alpha-diversity for some processes (55), such as nitrification. Moreover, increased evenness in the Z400 compared to Z50 biofilms could have resulted in lower abundance of specialized taxa (56, 57), such as *Nitrosomonas* and *Nitrospira*, and thereby decrease net NO_3_^-^ -production rates. Despite differences in relative abundance in *Nitrosomonas*, their absolute abundance was estimated to be the same in both Z50 and Z400. When measuring *Nitrosomonas* abundance, it is assumed that the entire population might contribute to aerobic ammonia oxidation. However we observed *Nitrosomonas* microcolonies throughout the Z400 biofilm depth (Fig 5A); the ones living in the deeper parts of the biofilm might have little or no access to oxygen (58). These *Nitrosomonas* cells could have low nitrification activity or they could represent strains capable of anaerobic respiration (59). Therefore, *Nitrosomonas* abundance might not be directly correlated with nitrification rates in thick biofilms.

A different ecosystem function, anaerobic NO_2_^-^ removal could occur via denitrification, anammox or DNRA. We observed higher anaerobic NO_2_^-^ removal rates in Z400 than Z50. This could be due to deterministic assembly, where the presence of anaerobic regions in Z400, likely allowed the establishment of taxa that could use NO_2_^-^ as electron acceptor. This agrees with a previous study (39), showing that an increase in biofilm thickness could lead to the emergence of new functions. Higher aerobic NO_2_^-^ removal in Z400 could occur because of NO_2_^-^ being used both as electron donor by NOB, and as electron acceptor by anaerobic taxa in anaerobic regions of the Z400.

In summary, we show that biofilm thickness can influence bacterial biofilm community composition despite the fact that history and all other external conditions are similar. The differences in communities between thin and thick biofilms were likely deterministic, but differences could not always be easily explained just by differences in redox conditions (*cf*. (30). Between-group beta-diversity was due to both nestedness and turnover, but dominated by turnover. Furthermore, based on potential and actual measurements, the two communities performed ecosystem functions at different rates, which support the idea that beta-diversity in the same metacommunity can lead to the emergence of multiple ecosystem functions (60). Results from these and similar experiments can be used in design of new process strategies in wastewater treatment. For example, thinner nitrifying biofilms could be combined with ticker biofilms to increase the number of ecological functions (39). Finally, bioreactors are well suited for experiments that can help disentangle factors of community assembly, as also suggested before (6).

## MATERIALS AND METHODS

### The reactor

The 0.5 m^3^ MBBR was located at the Sjölunda WWTP in Malmö, Sweden. The reactor was fed with effluent from a high-rate activated sludge process treating municipal wastewater (a feed with low carbon to nitrogen ratio). The average reactor load during one month before the sampling was 0.48 kg NH4+-N/m^3^,day and the NH4+ removal was 42%, at a pH of 7.4; dissolved oxygen (DO) concentration of 5 mg/L; and temperature of 17°C. After 261 days of operation, carriers were sampled for DNA-sequencing, FISH and activity tests to determine nitrogen transformations. The reactor contained a mixture of Z50 and Z400 carriers (Veolia Water Technologies AB – AnoxKaldnes, Lund, Sweden) at a total filling degree of approximately 30%. Thickness of the biofilm in Z-carriers is limited by a pre-defined grid wall height (36). Samples for optical coherence tomography measurements were taken on day 272 and data showed a biofilm thickness of 45 ±17 and 379 ± 47 (mean ± S.D.) µm for Z50 and Z400, respectively (37).

### Nitrogen transformation activity tests

Actual activity was measured in 1 L reactors in duplicate: Two reactors with 100 Z50 carriers each, and two with 100 Z400 carriers each. The incoming water was the same as the water feeding the 0.5 m^3^ reactor. At the time of measurement, the NH4+-N concentration was 19.6 mg/L, the DO was 5.5 mg/L, and the temperature was kept at 20°C. Mixing was achieved by supplying a gas mix consisting of N2-gas and air to the bottom of the reactors at an approximate total flow of 3 L/min and the DO was controlled to 5.5 mg/L by adjusting the amount of air in the gas mix. Nitrification rates were measured from mass balance as NO_2_^-^-N and NO_3_^-^-N mg/m^2^,day.

For the potential activity trials 3 L reactors, containing 400 carriers each, were used. The substrate consisted of NaHCO_3_^-^ buffer, pH adjusted to 7.5 using H_2_SO_4_, with phosphorous and trace minerals added in excess (36). Aerobic removal of NH_4_^+^ (starting concentration 35.2 NH_4_^+^- N mg/l) and NO_2_^-^ (starting concentration 32.5 NO_2_^-^-N mg/l) were measured separately in two different trials at 20°C for 1 hour, with sampling every 10 minutes. Mixing was achieved by supplying a gas mix consisting of N_2_-gas and air to the bottom of the reactors at an approximate total flow of 3 L/min. DO was controlled to 5.5 mg/L by adjusting the amount of air in the gas mix. Anaerobic trials of simultaneous removal of NH_4_^+^ and NO_2_^-^ (starting concentrations 35.5 NH_4_^+^-N and 36.1 NO_2_^-^-N mg/l) was measured at 30°C and were run for 2 hours with sampling every 20 minutes. Mixing was achieved by N_2_-gas from the reactor bottom. Before commencing the trials, the reactor with substrate was fed with N_2_-gas until the DO concentration was negligible and thereafter the carriers were added and the trials begun. Water samples were collected and filtered through 1.6 µm Munktell MG/A glass fiber filters and analyzed for NH_4_-N, NO_2_-N and NO_3_-N using standard Hach-Lange kits (LCK 303, 342 and 339, respectively).

### Fluorescence in situ hybridization (FISH)

FISH on cryosections and qFISH were done as previously described (37). The FISH probes used in this study are shown in Table S1. EPS and total nucleic acids on biofilm cryosections were stained with the FilmTracer SYPRO Ruby biofilm matrix stain and SYTO 40 (Thermo Fischer Scientific, USA), respectively. See Text S1, for details.

### Simulation of dissolved oxygen (DO) concentration profiles

A mathematical model was developed for simulating DO concentration profiles in the biofilms. The model considered the activities of aerobic heterotrophic bacteria, AOB and NOB. The bulk liquid concentrations of substrates (DO, nitrite, ammonium, and organic compounds), the measured biofilm densities, the microbial community compositions (as determined by FISH), the distribution of different functional groups of microorganisms in the biofilm (as measured by FISH), and kinetic coefficients from the scientific literature were used as input parameters. The thickness of the liquid boundary layer that limits diffusion of soluble substrates, including DO, from the bulk liquid to the biofilm was determined by comparing the ammonium oxidation rates calculated by the model to those measured during the nitrogen transformation activity tests. Since the exact concentrations of active biomass in the biofilms were unknown, the model was solved for different scenarios in which the active biomass was assumed to make up 20-80% of the measured total dry solids. It should be noted that the model only considers biofilm heterogeneity in one dimension (the depth direction). Layers parallel to the substratum are assumed to be homogenous. Real biofilms are three-dimensional structures containing channels and voids, which may allow oxygen transport into deeper regions locally. See Text S1 for details.

### DNA extraction and 16S rRNA gene sequencing

DNA was separately extracted from ten Z50 and ten Z400 carriers. DNA extraction, PCR and high throughput amplicon sequencing of 16S rRNA gene was done as previously described (61) with some modifications. Sequence variants (SVs) were generated for finer resolution of taxa (62, 63). See Text S1for details. Raw sequence reads were deposited at the NCBI Sequence Read Archive, no. SRP103666.

### Statistics

Data was analyzed in R (R Core Team 2018), using the packages Phyloseq (64), Vegan (65), DESeq2 (66) and betapart (42). Differential abundance of SVs was estimated with DESeq2 (66, 67), without random subsampling before the analysis. After independent filtering in DESeq2, 2578 of 3690 SVs were analyzed. A p_(adj)_ <0.01 value (DESeq2) was used as criterion for statistical significance. Subsampling to even depth was done prior to estimation of alpha-diversity and beta-diversity. Alpha-diversity was calculated as the first two Hill numbers (68), ^0^D (richness) and ^1^D (exponential of Shannon-Wiener index). Evenness was estimated as (^1^D/^0^D). Beta-diversity was estimated as pairwise Sørensen (β_sor_) dissimilarities, a presence-absence metric. Principal coordinate analysis (PCoA) was used for ordination. Permutational multivariate analysis of variance (PERMANOVA) (69) was used test for significant difference between group centroids. The components of β_sor_, turnover (β_sim_) and dissimilarity due to nestedness (β_sne_), were estimated as described by Baselga et al. (42) and used to calculate the beta diversity ratio (β_ratio_) as the ratio between β_sne_ and β_sor_ (44). If the β_ratio_ is smaller than 0.5, beta diversity is dominated by turnover rather than nestedness (44).

To disentangle the contribution of stochastic and deterministic community assembly mechanisms while at the same time accounting for possible differences in richness between Z50 and Z400, a null model approach was used. Firstly, the standardized effect size (SES) for pairwise Sørensen (SES_βsor_) dissimilarities were estimated in vegan using the oecosimu function. 999 null communities for estimation of SES_βsor_ were generated using the quasiswap algorithm (70), which preserve species richness and species incidence. For within groups null model analyses of Z50 and Z400 communities, only the taxa present in Z50 or Z400 respectively were used as the regional species pool. |SES_βsor_| > 2 was used as criteria to estimate if β_sor_ was different than expected by chance; a |SES| > 2 value is approximately a 95% confidence interval (71). Secondly, the RC_bray_ metric (72), which is based on quantitative data, was estimated for between-group comparisons, using 999 simulated communities. |RC_bray_| > 0.95 values were interpreted as deviations from the random expectation (47, 72).

## ACKNOWLEDGEMENTS

We thank Fred Sörensson for valuable discussions. The authors acknowledge the Genomics core facility at the University of Gothenburg, the Centre for Cellular Imaging at the University of Gothenburg and the National Microscopy Infrastructure, NMI (VR-RFI 2016-00968), for providing support and use of their equipment, and the colleagues at Veolia Water Technologies AB – AnoxKaldnes, Lund, Sweden, for monitoring the pilot reactor. This work was funded by FORMAS (Contract no. 243-2010-2259, 211-2010-140, 2015-1515-30425-28, 245-2014-1528, 942-2015-683 and 2012-1433), SVU (Contract no. 10-105), the Foundations of Carl Trygger (CTS 12:374), Adlerbertska forskningsstiftelsen, Wilhelm & Martina Lundgrens Vetenskapsfond (2015-0317, 2015-0309) and the Swedish Water & Wastewater Association via the research cluster VA-teknik Södra.

## Conflict of interest

The authors declare no conflict of interest, MP work at Veolia Water Technologies AB – AnoxKaldnes, Lund, Sweden.

## SUPPLEMENTAL INFORMATION

**Fig. S1. Log2fold (DESeq2) changes for the 40 most abundant SVs based on average abundance**

Phylum, order and genus classification are shown. Each circle represents an SV. The size of the circle is proportional to the total sequence read abundance for the SV. A negative log2 fold change indicates that SV are more abundant in Z400 biofilm, while a positive log2 fold change indicates SVs more abundant in Z50 biofilms.

**Fig. S2. Log2fold (DESeq2) changes for *Bdellovibrionales* SVs.**

Genus classification is shown. Each circle represents an SV. The size of the circle is proportional to the total sequence read abundance for the SV. A negative log2 fold change indicates that SV are more abundant in Z400 biofilm, while a positive log2 fold change indicates SVs more abundant in Z50 biofilms. SVs with a NA p_(adj)_ value (DESeq2) are not shown.

**Fig. S3. Density profiles and biomass distribution for the DO model**

A and B: Biofilm density profiles (total dry solids) in the Z50 and Z400 biofilms respectively. C: Assumed biomass distribution in the Z400 biofilm based on input from qFISH and cryosection FISH images

**Table S1.** FISH probes used in this study

**Table S2**. Notation used in the DO model.

**Table S3.** Kinetic rate expressions used in the DO model.

**Table S4.** Stoichiometric matrix used in the DO model

**Table S5**. Kinetic and stoichiometric coefficients used in the DO model.

**Table S6**. Default input values for physical parameters used in the DO model.

**Text S1.** Supplemental Material and Methods

